# Neurodevelopmental disease genes implicated by *de novo* mutation and CNV morbidity

**DOI:** 10.1101/209908

**Authors:** Bradley P. Coe, Holly A.F. Stessman, Arvis Sulovari, Madeleine Geisheker, Fereydoun Hormozdiari, Evan E. Eichler

## Abstract

We combined *de novo* mutation (DNM) data from 10,927 cases of developmental delay and autism to identify 301 candidate neurodevelopmental disease genes showing an excess of missense and/or likely gene-disruptive (LGD) mutations. 164 genes were predicted by two different DNM models, including 116 genes with an excess of LGD mutations. Among the 301 genes, 76% show DNM in both autism and intellectual disability/developmental delay cohorts where they occur in 10.3% and 28.4% of the cases, respectively. Intersecting these results with copy number variation (CNV) morbidity data identifies a significant enrichment for the intersection of our gene set and genomic disorder regions (36/301, LR+ 2.53, p=0.0005). This analysis confirms many recurrent LGD genes and CNV deletion syndromes (e.g., *KANSL1, PAFAH1B1, RA1,* etc.), consistent with a model of haploinsufficiency. We also identify genes with an excess of missense DNMs overlapping deletion syndromes (e.g., *KIF1A* and the 2q37 deletion) as well as duplication syndromes, such as recurrent *MAPK3* missense mutations within the chromosome 16p11.2 duplication, recurrent *CHD4* missense DNMs in the 12p13 duplication region, and recurrent *WDFY4* missense DNMs in the 10q11.23 duplication region. Finally, we also identify pathogenic CNVs overlapping more than one recurrently mutated gene (e.g., Sotos and Kleefstra syndromes) raising the possibility that multiple gene-dosage imbalances may contribute to phenotypic complexity of these disorders. Network analyses of genes showing an excess of DNMs confirm previous well-known enrichments but also highlight new functional networks, including cell-specific enrichments in the D1+ and D2+ spiny neurons of the striatum for both recurrently mutated genes and genes where missense mutations cluster.

## INTRODUCTION

The importance of de *novo* mutations (DNMs) underlying neurodevelopmental disorders (NDDs) has been recognized for many years. Some of the strongest genome-wide evidence came from early copy number variation (CNV) studies, which consistently showed an excess of de novo as well as large private CNVs in patients with autism, developmental delay and epilepsy^1–4^. Because of the interspersed duplication architecture of the human genome^5^, evidence of recurrent mutation in probands was often readily identified among a few hundred families—at least for those CNVs flanked by segmental duplications (SDs)^2^ or mapping near telomeres^6^. Significance based on recurrence was more readily achieved from smaller sample sizes because of elevated CNV mutation rates in these specific regions of the genome, although in most cases the individual genes underlying the genomic disorders remained unknown.

The advent of next-generation sequencing and exome sequencing rapidly accelerated our ability to specify genes associated with potentially pathogenic de novo single-nucleotide variants (SNVs) for both developmental delay and autism^7,8^. The locus heterogeneity of these pediatric disorders, and the more random nature of SNV mutations, however, has made it more difficult for individual studies to provide statistical significance for DNM recurrence^9,10^. Moreover, different statistical models for discovery of genes with potentially pathogenic DNMs have been developed, including those based on chimpanzee–human divergence, trinucleotide mutation context, and clustering of DNMs ^13–15^. Finally, despite extensive CNV analyses of nearly 45,000 patients with autism and developmental delay^16–18^, few attempts have been made^18^ to integrate the wealth of CNV data with recent exome sequencing results despite the fact that the same model of dosage imbalance might underlie a common genetic etiology for a specific locus (i.e., likely gene-disruptive (LGD) mutations and CNV deletions).

In this study, we perform an integrated meta-analysis of DNMs that have been recently published and compiled based on exome sequencing data of 10,927 individuals with a diagnosis of autism spectrum disorder (ASD) or intellectual disability and/or developmental delay (ID/DD)^19^. We overlay these data with known genomic disorders and a CNV morbidity map generated from 29,085 cases and 19,584 controls and jointly consider data from parent–child trios from ASD and DD cohorts in order to increase statistical power. This treatment of the data is justified because of the significant comorbidity between these two disorders^20,21^ and because autism cases with a severe DNM are enriched in DD^22^. We compare two statistical models of recurrence as well as mutational classes because of potentially different functional consequences (e.g., recurrent missense vs. LGD DNMs). The goals of this study are threefold: 1) provide an integrated list of candidate NDD genes based on multiple lines of DNM and CNV evidence, 2) compare different models of recurrent mutation, and 3) identify the most likely genes underlying pathogenic microdeletion and microduplication CNVs associated with DD.

## RESULTS

### Genes enriched for de novo SNV mutation and model comparisons

We compiled de *novo* variation identified from exome sequencing of 10,927 cases with NDDs from the database denovo-db v. 1.5 release^19^. This includes 5,624 cases with a primary diagnosis of ASD and 5,303 cases with a diagnosis of ID/DD collected from 17 studies ^11,22–37^. We considered all protein-altering and LGD mutations, including frameshifts, splice donor/acceptor mutations, start losses and stop gains. The combined set of 12,172 DNMs included 2,357 LGD and 9,815 missense mutations.

We initially applied two statistical models. The first incorporates locus-specific transition/transversion/indel rates and chimpanzee–human coding sequence divergence^11^ to estimate the number of expected DNMs, hereafter referred to as the chimpanzee–human divergence model or CH model. The second model, denovolyzeR, estimates mutation rates based using trinucleotide context and incorporates exome depth and divergence adjustments based on macaque–human comparisons over a +/-1 Mbp window and accommodates known mutational biases such as CpG hotspots. Both models apply their underlying mutation rate estimates to generate prior probabilities for observing a specific number and class of mutations for a given gene. While both models incorporate LGD and missense probabilities, we recently modified the CH model to incorporate CADD scores^38,39^ allowing us to also specifically test for enrichment for the most severe 0.1% missense mutations (i.e., CADD scores over 30 or MIS30). Such missense mutations are more likely to be functionally equivalent to an LGD mutation and have been shown to be significantly enriched in NDD cases compared to controls^14^. Each analysis was corrected genome-wide for the number of genes present in the corresponding models (18,946 for CH model and 19,618 for denovolyzeR). Based on prior assumptions that 50% of DNMs will likely be disease related, we have chosen a q-value cutoff of 0.1 for both models to represent a false discovery rate (FDR) near ~5%^18,40^. We note that a more stringent q-value threshold of 0.05 slightly reduces the number of significant genes. Only 16% (24/147) of the genes which show an excess of de novo missense mutations demonstrate a q-value between 0.05 and 0.1 (four of which are significant by LGD or MIS30 DNM).

Combined, the two models (union set) identified 301 candidate NDD genes with evidence of excess DNM with a q-value < 0.1, and at least two mutations for at least one mutational category (Table 1, Table 2, Supplemental Table 1). This includes 163 genes with excess LGD mutations and 147 genes with excess missense mutations. Among these, 31 genes demonstrate evidence of both LGD and missense mutations (Figure 1, Table 1, and Supplemental Table 1). In general, both models highlight similar genes (Figure 1), particularly for LGD events where 71.2% (116/163) of genes are shared. This stands in contrast to recurrent missense DNMs where only 47.6% (70/147) of the genes overlap between the models. With a few exceptions (Figure 1C-D, Table 1), genes unique to a model are near the boundary of statistical significance. The most substantial outliers are unique to the CH model (LGD outliers: *NONO, MEIS2, LEO1, WDR26, CAPRIN1*; missense outliers: *CAPN15, SNAPC5, DLX3, TMEM178A, ADAP1, SNX5, SMARCD1, WDR26, AGO)* and in all cases represent genes with no variation in the coding sequence between the chimpanzee and human reference sequences used to build the model.

**Table 1:** Recurrent DNM gene summary and model comparison.

| Variant Category | CH Count | denovolyzerR<br>Count | Union<br>Count | Intersection<br>Count | ID/DD Only | ASD Only |
| --- | --- | --- | --- | --- | --- | --- |
|  |  |  |  |  | Union<br>Count | Union<br>Count |
| LGD & Missense_30 & Missense | 14 | NA | 14 | NA | 4 | 0 |
| LGD & Missense_30 | 1 | NA | 1 | NA | 0 | 0 |
| LGD & Missense | 14 | 24 | 16 | 22 | 22 | 4 |
| LGD | 104 | 122 | 132 | 94 | 103 | 35 |
| Missense_30 & Missense | 32 | NA | 34 | NA | 11 | 0 |
| Missense_30 | 23 | NA | 21 | NA | 0 | 0 |
| Missense | 58 | 75 | 83 | 48 | 80 | 6 |
| *total* | 246 | 221 | 301 | 164 | 220 | 45 |

**Table 2:**
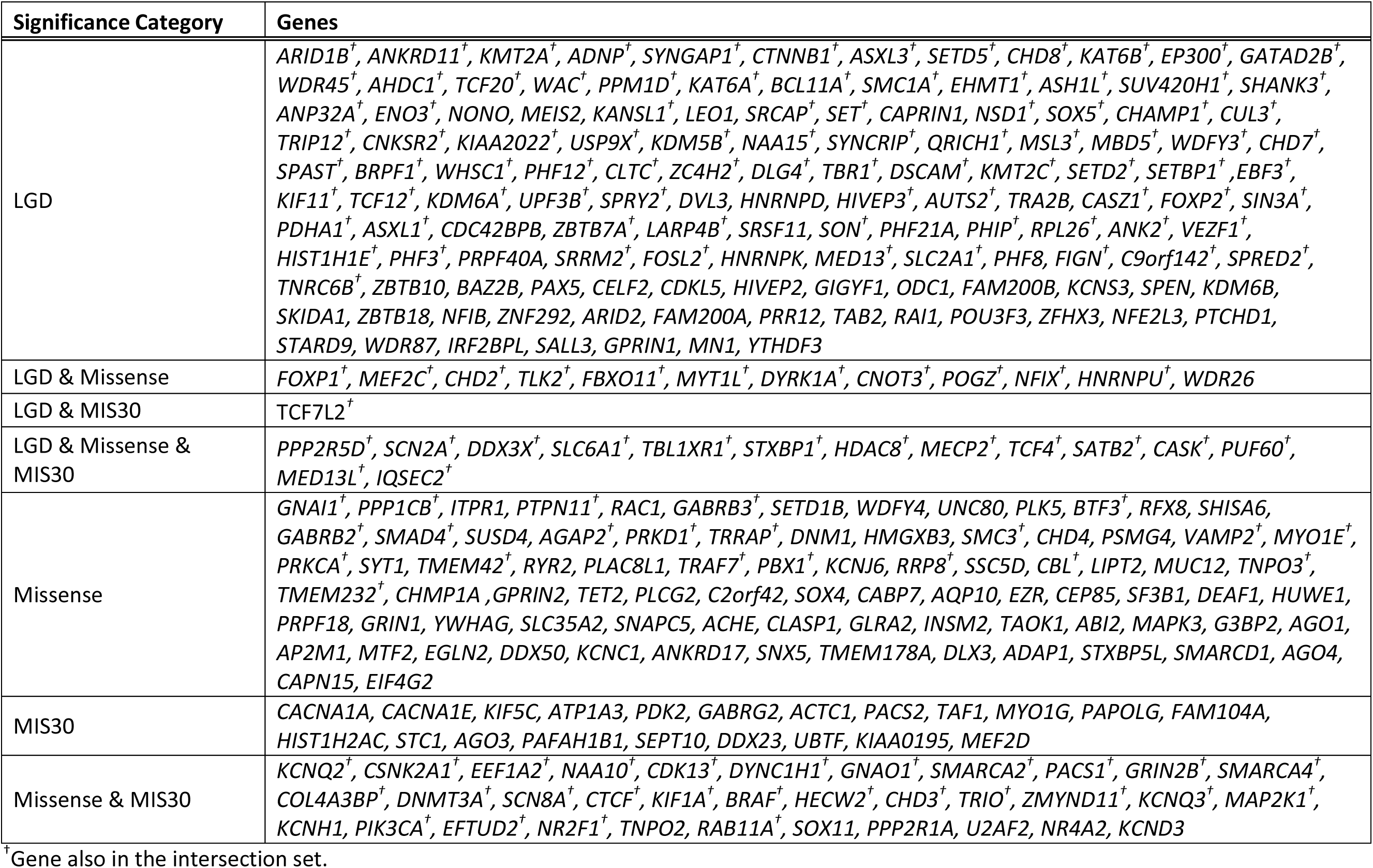
Genes enriched for *de novo* variation in 10,927 ASD/ID/DD patients in denovo-db v1.5

**Figure 1:**
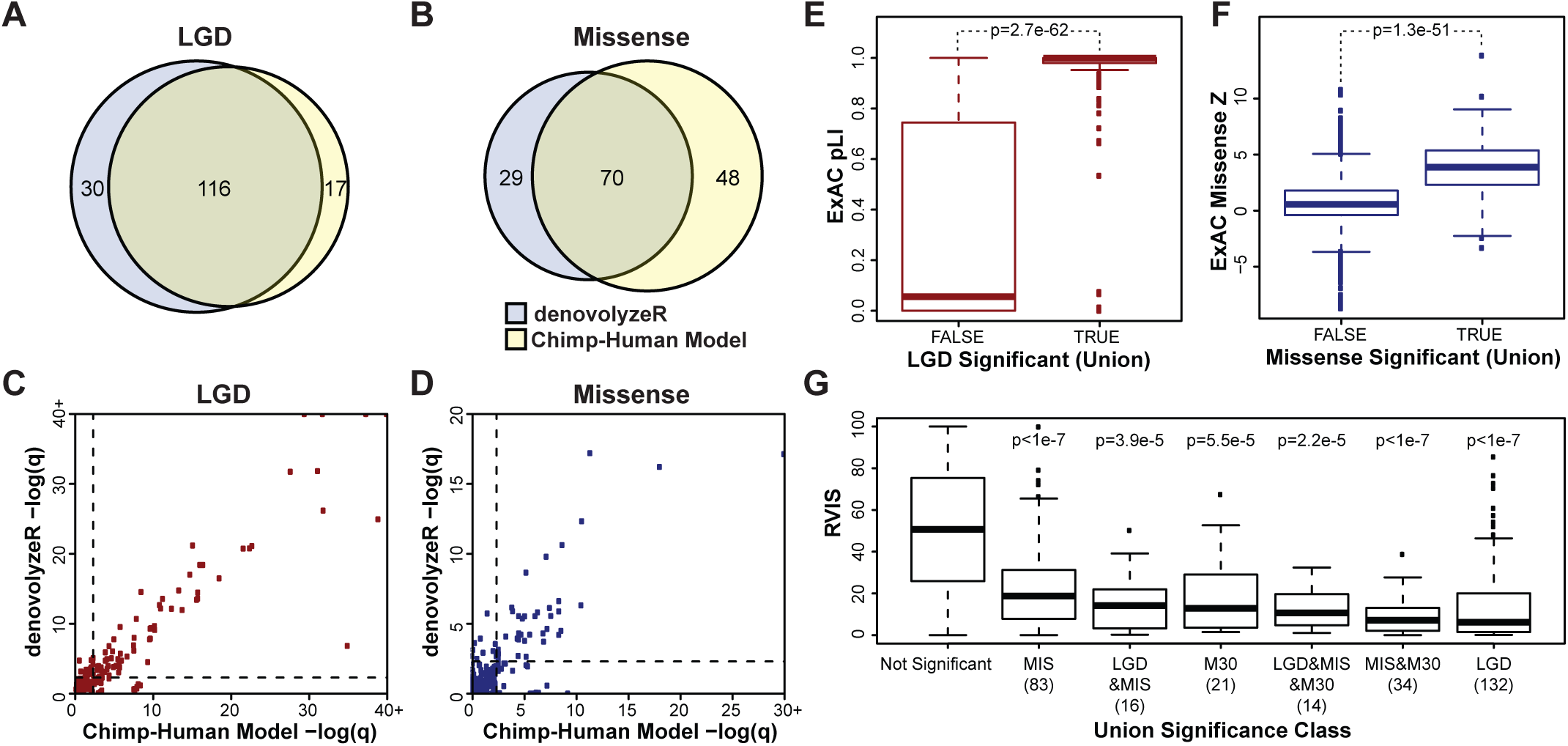
*de novo* enriched genes and their characteristics. Shown are the results of applying both the chimpanzee-human (CH) divergence model and denovolyzeR to de novo variation in 10,927 individuals with ASD/ID/DD. The two models show considerable gene overlap **(A,B)** with correlated significance values (LGD r^2^ = 0.94, missense r^2^ = 0.74) **(C,D)**. CH model LGD outliers include *NONO, MEIS2, LEO1, WDR26,* and *CAPRIN1,* and denovolyzeR LGD outliers include *ZBTB18* and *FAM200B***(C)**. CH model missense outliers include *CAPN15, SNAPC5, DLX3, TMEM178A, ADAP1, SNX5, SMARCD1, WDR26,*and *AGO4,* and denovolyzeR missense outliers include *ITPR1, RAC1, SETD1B, WDFY4,* and *UNC80* **(D)**. Recurrent mutated LGD genes (true) are highly enriched for genes intolerant to mutation as defined by ExAC pLI score **(E)**. Genes significantly enriched for missense DNMs are outliers by the ExAC missense depletion Z scores **(F)**. Similarly, all subcategories of significant genes are intolerant to mutation (RVIS percentile) when compared to non-significant genes (p-values are corrected for all possible group comparisons). The LGD and MIS30 group is not shown as it only represents one gene *(TCF7L2,* RVIS percentile = 10.96) **(G)**.

Since it has been well-established that NDD genes are less tolerant to mutation, we categorized the 301 genes into different functional groups and compared their tolerance to mutation in the general population using three metrics: residual variance to intolerance score (RVIS); probability of loss-of-function intolerance (pLI); and missense constraint scores (missense Z scores). For both pLI and missense Z scores, we utilized the ExAC subset with known neuropsychiatric cohorts removed (45,376 individuals)^41^. For genes with an enrichment of LGD variants, we observe a significant increase in pLI scores compared to all other genes, which is almost predictive (p = 2.7e-62, ROC AUC = 0.89) (Figure 1E). We also observed a significant increase in missense constraint (missense Z scores) among genes with enrichment for missense variation (p = 1.3e-51, ROC AUC = 0.84) (Figure 1F). Using RVIS as a constraint metric demonstrates a similar trend. We observe a significant RVIS depletion for all categories where at least two genes were identified (Figure 1G). Examination of a combination of constraint metrics is particularly valuable as a small number of genes demonstrate conflicting results, such as the LGD- and missense-enriched gene *MEF2C,* which is involved in severe ID when disrupted by deletions or mutations^42^ and demonstrates constraint by RVIS, but not by pLI (RVIS = 18.97, pLI = 2.4e-3, missense Z score = 4.47).

A small number of genes are enriched for recurrent DNMs but do not demonstrate conservation by either RVIS or pLI/missense Z scores. For example, there are 14 genes significant for LGD recurrence with a pLI below 0.9 and RVIS percentile above 20 and these would be predicted to be enriched for false disease associations^43^. However, half (7/14) of these genes have an established association with ASD or ID/DD, including four X chromosome-linked genes *(MECP2, ZC4H2, UPF3B, HDAC8)^44^–^41^* and three autosomal genes *(ASXL1, PURA,* PPM1D)^48-50^. The remaining seven genes lack support in the literature *(ENO3, HIVEP3, C9orf142, KCNS3, NFE2L3, SALL3, GPRIN1)* strongly linking them to ID/DD or ASD. In addition we should note that a subset of genes show evidence of DNM in controls. Among the 2,278 controls in denovo-db v1.5 82.7% (249/301) of the union genes have no detected LGD or Missense DNM in controls. Among the 52 genes with at least 1control DNM, 14 genes have control DNM matching the primary significance category including 3 LGD in *KDM5B* and 13 genes with a single control DNM (1 LGD in *KCNS3, TCF12;* 1 Missense in *AGAP2, AGO4, CAPN15, GLRA2, GRIN1, PBX1, SF3B1, WDFY4, TNPO3;* and 1 MIS30 in *CACNA1E, MEF2D).* The detection of control events may represent incomplete penetrance, variable expressivity, undiagnosed/subclinical controls, or benign variation (primarily in the case of missense variation). None of the recurrent control DNM genes reach exome wide FDR significance. While some of these are plausible candidates, disease significance should be considered with caution until additional functional and clinical data establish their role.

#### ASD versus DD genes

Because the number of samples ascertained for ASD and DD are approximately equal, we examined the distribution of LGD and missense mutations between the two populations. The majority of the genes—76% (229/301) of the union and 84% (137/164) of the intersection—show evidence of DNM in both ASD and ID/DD cohorts. A small number of genes are specific or enriched. For example, we directly compared the variant distributions among the genes with DNM and identified four genes *(ANKRD11, KMT2A, ARID1B, DDX3X)* with recurrent LGD and one gene *(KCNQ2)* with recurrent missense mutations as being strongly biased toward a diagnosis of ID/DD (q < 0.1). Although no genes are statistically enriched for ASD (Figure 2), several genes (e.g., *WDFY3,* nominal p = 0.019; *DSCAM,* nominal p = 0.036; *CHD8,* nominal p = 0.074) tend toward ASD diagnosis. While larger sample sizes are still required, these results highlight the significant comorbidity of ID/DD and ASD but also suggest that there will be genes that associate preferentially with ID or ASD. Considering all 301 genes and the full set of NDD patients, we calculate that 19.1% (2,085/10,927) of samples have at least one *de novo* event in this gene set. The proportion of patients with a DNM is significantly higher for ID/DD (28.4% or 1,507/5,303 patients) when compared to ASD (10.3% or 578/5,624 patients) (OR = 3.47, p = 1.22e-131).

**Figure 2:**
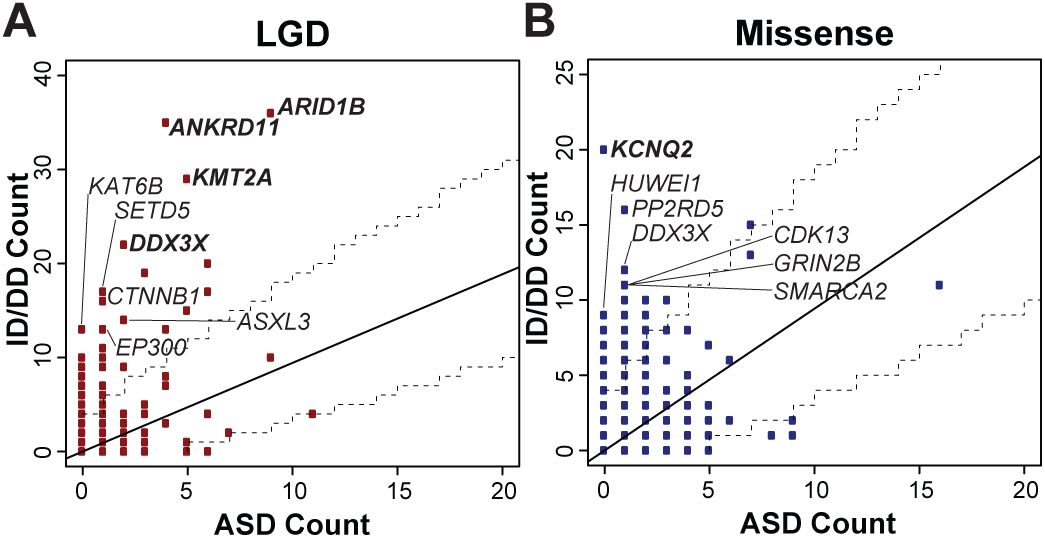
Comparison of de *novo* variation rates in ASD and ID/DD. The plots compare DNM rates for genes for patients from ASD and ID/DD studies included in our combined analysis. More than 75% of genes show DNM in both ASD and DD patients. We identify four LGD genes *(ARID1B, ANDKRD11, KMT2A, DDX3X)***(A)** and one missense gene *(KCNQ2)***(B)** that are biased for an ID/DD diagnosis at a q- value threshold of 0.1. Additional candidates for phenotypic bias at nominal significance (dashed lines at p = 0.05) were also identified. Larger cohorts will be needed to confirm gene biases, especially with respect to ASD.

#### Network enrichment

Examination of the 301 genes identified by the union of both statistical approaches suggests that our set is strongly enriched for functionally related networks of genes. The STRING database, for example, identifies a highly significant 1.8-fold (1,283 edges vs. 699 expected) enrichment in interactions among the 301 union genes (p < 2.2e-16). Given this high level of interconnectivity, we applied MAGI^51^, a gene network discovery tool, to identify potential gene clusters, functional enrichments, and additional candidate interactions. Here, we present the top four networks from this analysis and their associated PANTHER functional enrichments (Figure 3A-D). Module 1 (20 genes) highlights “regulation of transcription from RNA polymerase II promoter” (p = 0.0269) (Figure 3A) and contains 15 significant genes in addition to three additional candidates that do not yet reach significance *(CREB1, RBBP5, CBX5)* and two genes (*SREK1*, *SMARCB1)* with no DNM in our current data set. Module 2 highlights multiple functions relating to neurotransmitter signaling (p = 0.0358) and synaptic signaling (p = 8.91e-5) (Figure 3B) and contains 11 significant genes in addition to eight genes that do not reach significance *(DLG2, HTT, AP2A2, KCNJ4, KCNB1, STX1A, GRIN2A, CAMK2A)* and one gene with no DNM *(PRKCB).* Module 3 highlights the “transmembrane receptor protein serine/threonine kinase signaling pathway” (p = 0.002) (Figure 3C) and contains nine significant genes in addition to 20 genes that do not reach significance *(SMURF2, SMURF1, CDC73, RNPS1, RBBP4, UBE3A, CUL1, FBXW11, VCP, VPS4A, PPP5C, PRPF38A, SKIL, HSPA4, PSMD3, UIMC1, GAPVD1, NLGN2, GTF3C1, NRXN1),* and six genes with no DMN *(ING2, SRSF4, FAF1, UBC, HSP90AB1, YWHAB).* Finally, Module 4 highlights c-Jun N-terminal kinase (JNK) (p = 4.65e-5) and Mitogen-activated protein kinase (MAPK) (p = 3.13e-6) cascades (Figure 3D) and contains two significant genes in addition to 15 genes that do not reach significance *(RPS6KA3, RASGRF1, MAPK8IP1, SMAD3, DUSP3, MAPK9, SPTBN1, ACTN4, CAMK2G, TFE3, PRKAR1A, SNAP25, MAPK8IP2, MAPK8IP3, PRKAR1B)* and three genes with no DNM *(SYN1, MAPK1, PRKACA).* Among these nonsignificant genes are many previously identified NDD candidate genes (e.g., *NRXN1, GRIN2A,* CAMK2A)^52–54^ suggesting this group as a potential target for future screening and disease gene discovery.

**Figure 3:**
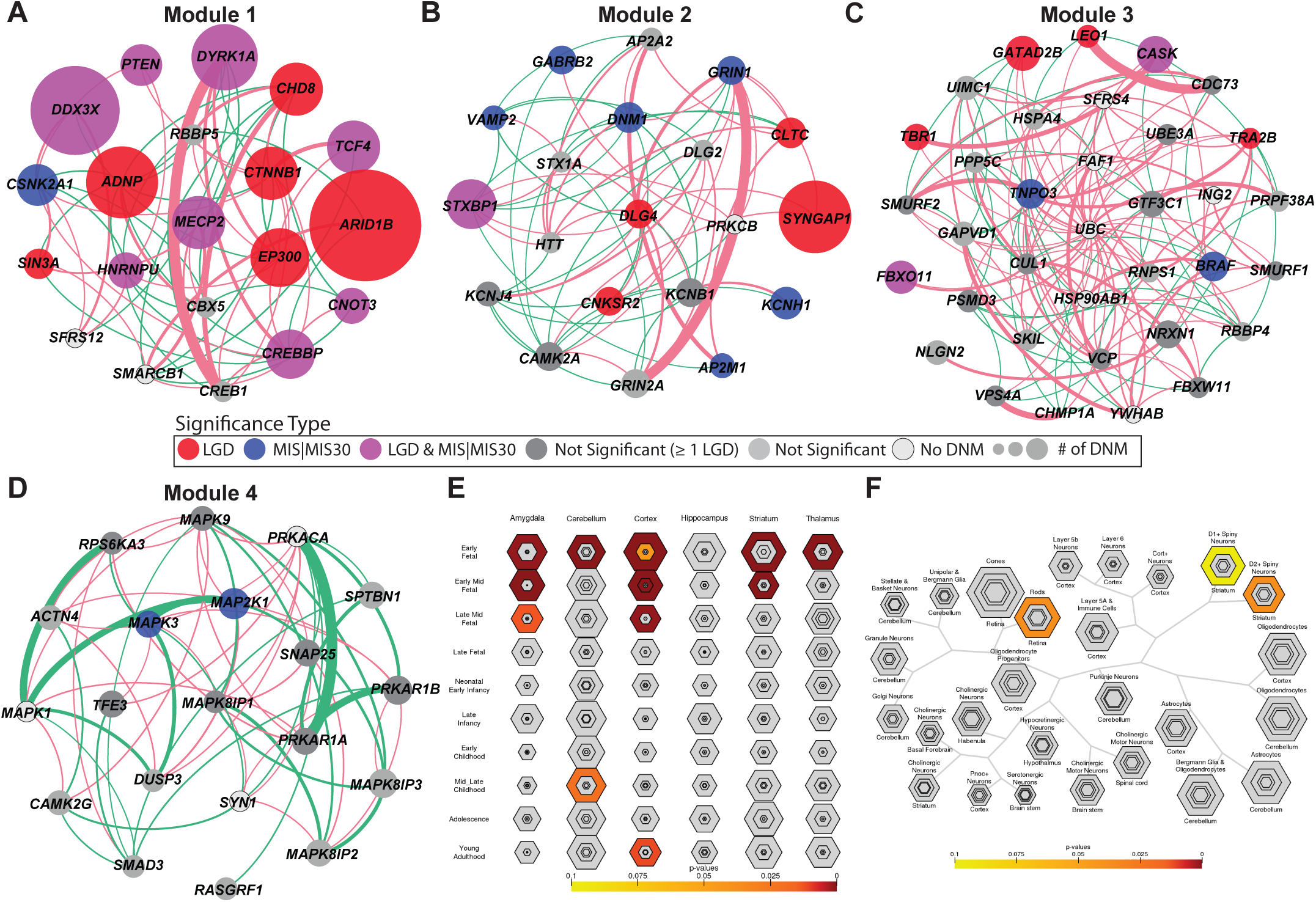
Gene expression and protein-interaction networks. The plots compare DNM rates for genes for patients from ASD and ID/DD studies included in our combined analysis. More than 75% of genes show DNM in both ASD and DD patients. We identify four LGD genes *(ARID1B, ANDKRD11, KMT2A, DDX3X)***(A)** and one missense gene *(KCNQ2)***(B)** that are biased for an ID/DD diagnosis at a q- value threshold of 0.1. Additional candidates for phenotypic bias at nominal significance (dashed lines at p = 0.05) were also identified. Larger cohorts will be needed to confirm gene biases, especially with respect to ASD.

In addition to enrichment in the PPI context, we also note strong functional cell-specific and tissue-specific enrichment analyses (CSEA and TSEA)^55,56^. As expected, the 301 gene set is enriched for the brain expression with a bias toward early to mid-fetal gene expression in the cortex, striatum and amygdala (Figure 3E). Among these the greatest specificity is observed for early to early mid-fetal cortical development. By CSEA, the *de novo* gene set shows enrichment in both classes of medium spiny neurons within the striatum (striatum D1+ and D2+ medium spiny neurons BH corrected p = 0.043 and p = 0.02) at a pSI (specificity index p-value) threshold of 0.05. Additionally, we observe nominal significance for D1+ and D2+ spiny neurons (uncorrected p = 0.018 and p = 0.015) at a pSI of 0.01 (Figure 3F). This cell-specific enrichment appears to be driven by genes with an excess of LGD DNMs (n = 163) (BH p = 0.029 for D1+ and D2+ neurons at pSI = 0.05, nominal p = 0.009 and 0.008 at pSI = 0.01), but not among genes enriched for recurrent missense mutations.

#### Clustered missense

As a final form of candidate gene annotation, we examined the distribution of missense DNMs for potential clustering within the predicted protein as previously described^14^. We identified a set of 183 nominally significant genes (using ExAC as a reference). Interestingly, 78% of this gene set does not overlap genes that reach *de novo* significance for missense recurrence (Supplemental Table 1). Examination of this clustered missense set by CSEA once again highlights the D1+ and D2+ spiny neurons of the striatum (nominal p = 0.066 and p = 0.005, respectively, at pSI = 0.05). Several D1+ genes were identified as significant for recurrent missense and clustered mutations *(PPP2R5D, SMARCD1, GRIN2B).* Among those showing evidence of clustered DNM exclusively, we identify genes enriched for D1+ *(PPP2R5D ARID1A, CEP131, INF2, RYR3, PHACTR1, KDM3A, GRIN2B, SLITRK5)* and D2+ *(ARID1A, CEP131, INF2, RYR3, AFF3, MAPKBP1, PHACTR1, KDM3A, MYO18A, SMAD4, SLITRK5)* spiny neuron expression. Mutations in these genes should be considered as candidates for further follow-up since not all genes reach exome-wide statistical significance. However, 99 genes remain significant after multiple testing corrections at a q-value threshold of 0.05. This set is significant for gene enrichment only in D2+ *(PRKD1, ARID1A, INF2, RYR3, AFF3, MAPKBP1, SLITRK5, KDM3A, MYO18A)* (BH p = 0.002) but not in D1+ (BH p = 0.112) spiny neurons.

#### Projected rates of gene discovery

Based on the number of genes that reach significance for DNM in our cohort of 10,927 cases, we estimated the potential yield by mutational class and the CH model. To this end, we subsampled smaller populations from our set 10,000 times each and tested for how many genes would reach significance using the CH model in a resampled cohort of similar phenotypic composition (i.e., an NDD cohort composed of a nearly equal number of patients with autism and ID/DD). We assessed logistic growth models for each mutation class and selected the best fitting model by Bayesian information criteria (BIC) to predict future performance. For genes with excess LGD DNMs, we observe what appears to be a rapid upcoming plateau in gene discovery with an asymptote at 213 genes (95% CI 203-225) (ΔBIC linear model - Weibull model = 402) (Figure 4). Similarly, for genes with an excess of MIS30 DNMs the model predicts an asymptote of only 100 genes (95% CI 95-105) (ΔBIC linear model - Weibull model = 235) (Figure 4). By contrast, genes with an excess of recurrent missense mutations cannot yet be projected. The data does not fit any logistic growth model and only poorly fits a linear model as the initial part of the curve demonstrates expected exponential growth. Further parsing the missense signal by examining missense mutations with CADD scores under 30 by a modified CH model also fails to generate a predictable asymptote suggesting that this category will represent the largest yield for novel genes as new exomes are sequenced.

**Figure 4:**
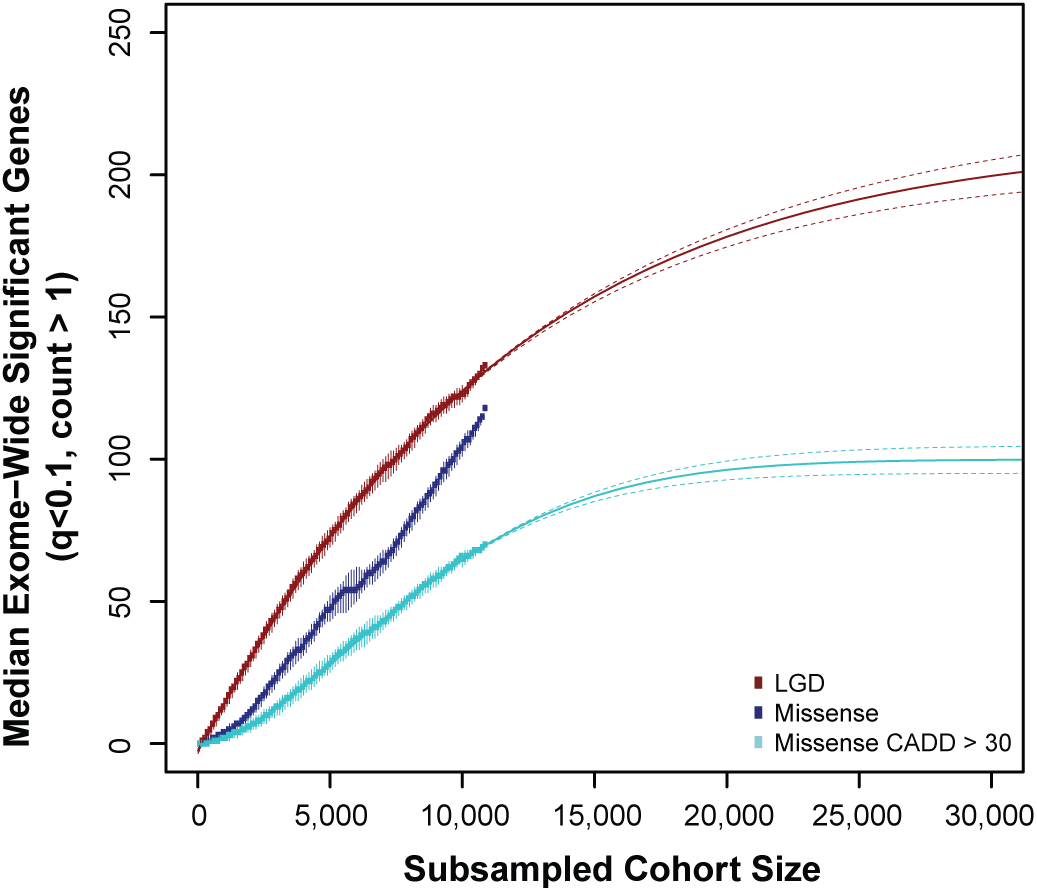
Estimation of gene discovery rates in future cohorts. We estimate the number of genes reaching significance under the CH model at varying population sizes subsampled from the total cohort of 10,927 individuals. Both the number of significant genes with recurrent LGD and missense (CADD score >30) DNM appear to be saturating with limited new gene discovery as sample sizes grow. De *novo* missense variants (including MIS30), however, as a more general class demonstrate a more complex growth pattern with no best-fit line and, thus, likely represents the most important reservoir for new gene discovery as sequence data are generated from additional ASD and DD cohorts.

#### CNVintersection

In order to identify potentially dosage-sensitive genes underlying pathogenic CNVs, we intersected the 301 candidate gene set with a list of 58 genomic disorders based on previous CNV morbidity maps and the DECIPHER database (Table 3, Figure 5). Considering all genes with a *de novo* variant (n = 6,886), we find that 36 of 301 significant genes intersect a genomic disorder region. This represents a significant (p = 0.0005) enrichment (LR+ 2.53 [95% CI 1.43 - 1.91]) compared to expectations supporting the notion that CNVs and DNMs converge on a common genetic etiology of gene-dosage imbalance.

**Table 3:**
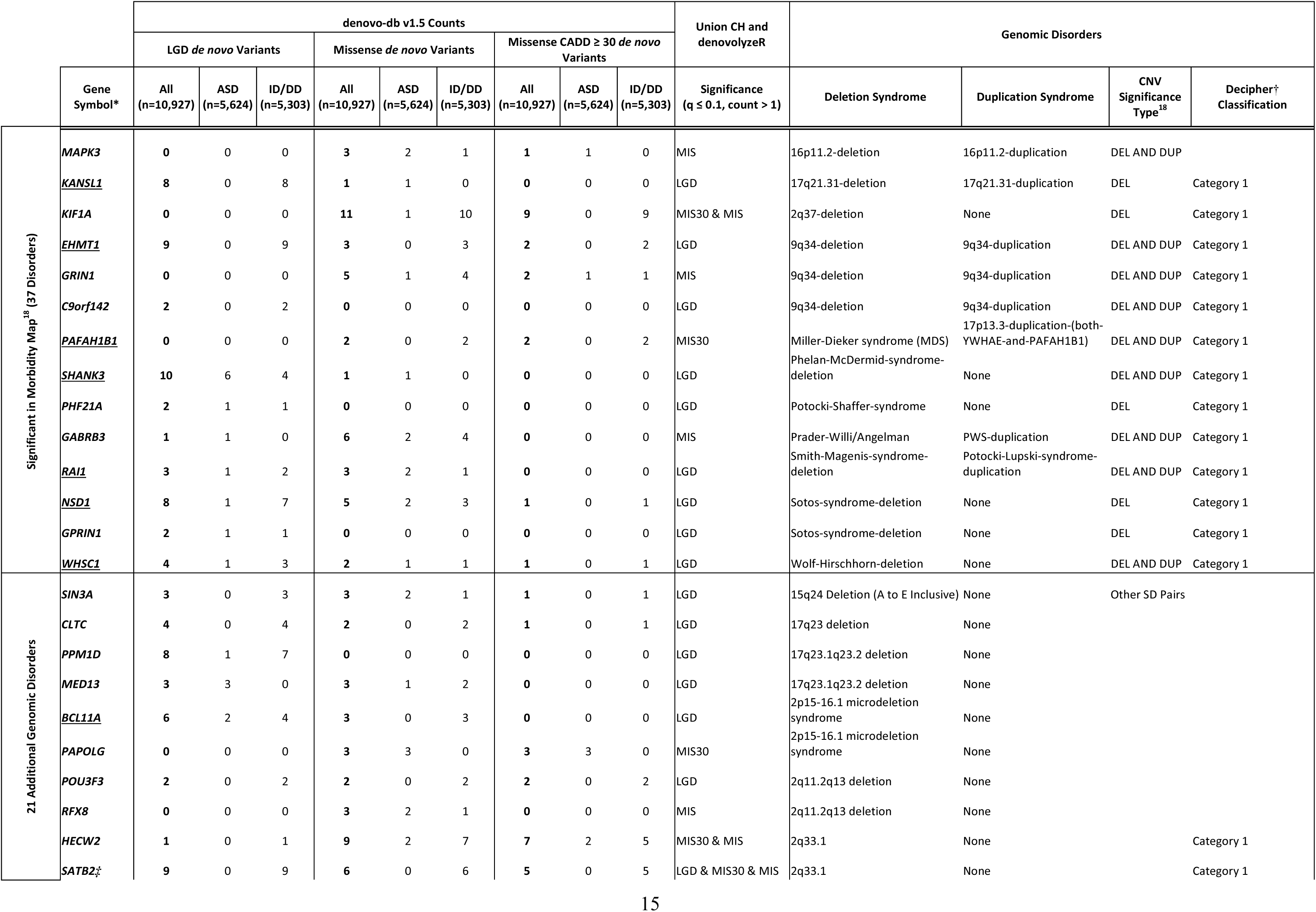
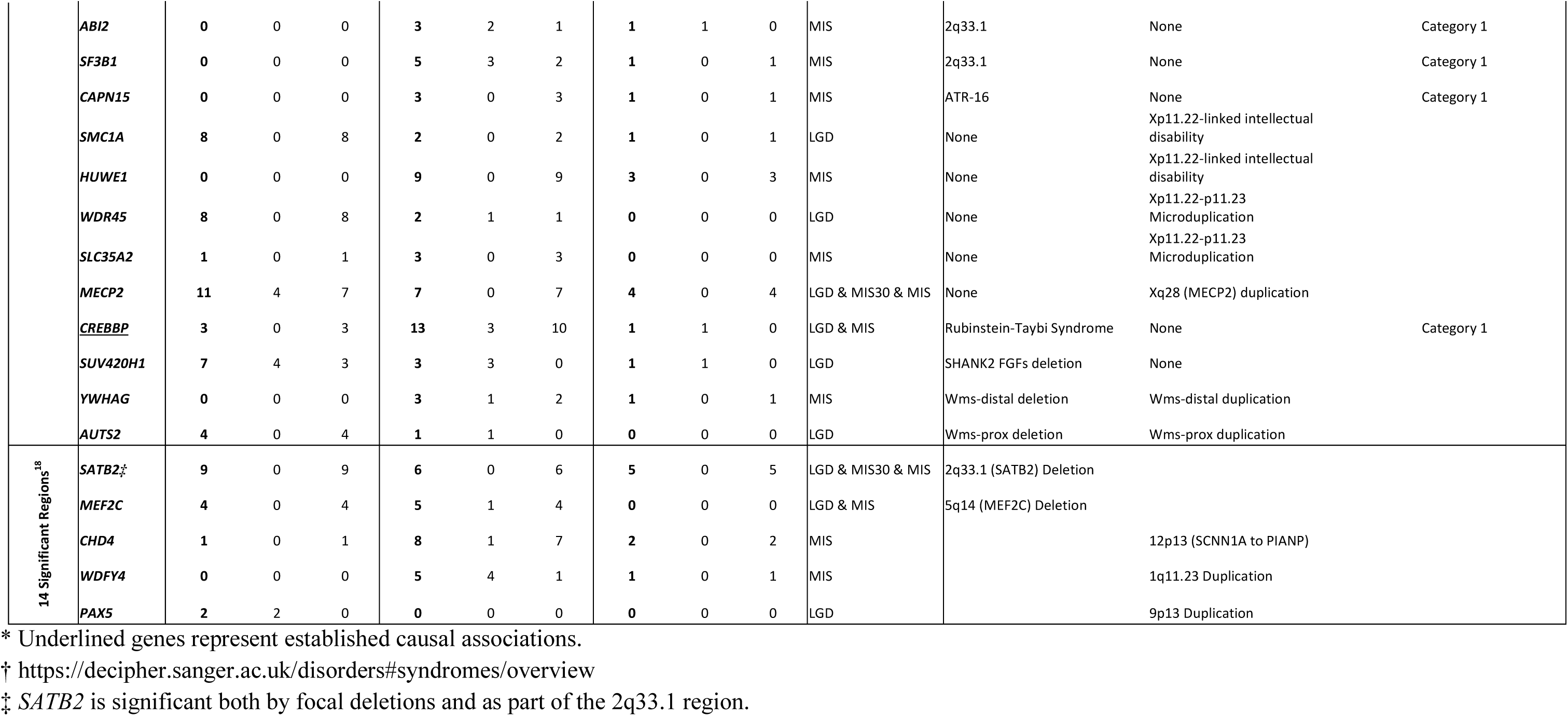
Intersection between pathogenic CNVs and recurrently mutated genes

**Figure 5:**
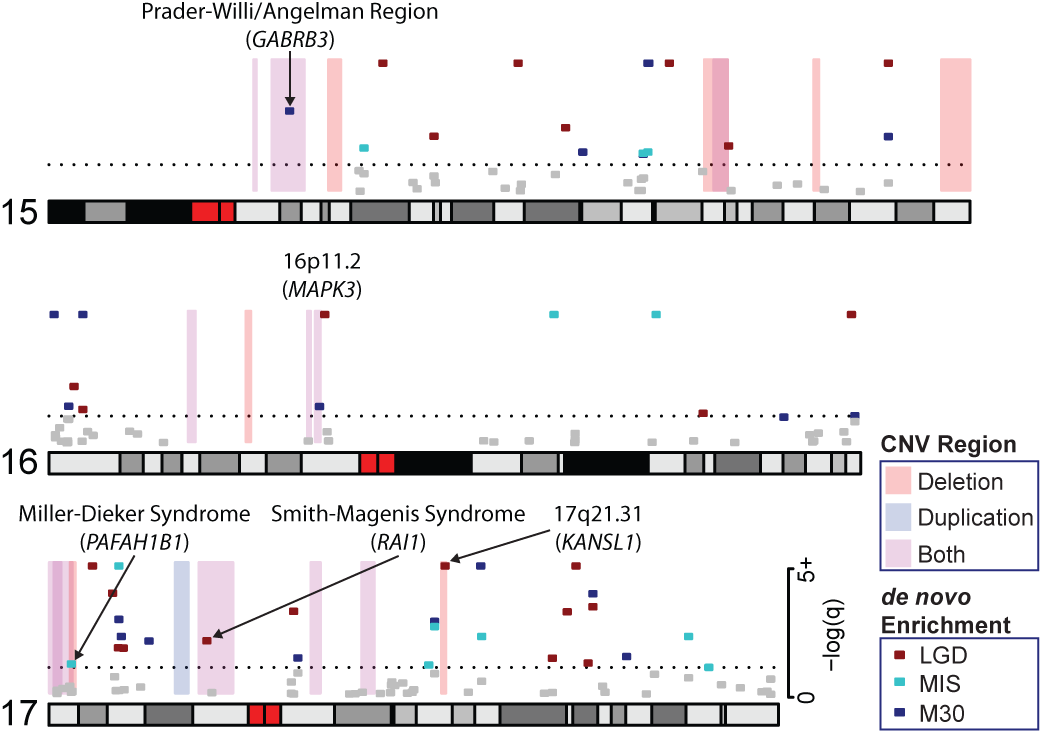
Integration of *de novo* SNVs and CNV morbidity map. Shown are examples of pathogenic CNVs (blue, red and purple shading) associated with genomic disorders from chromosomes 15, 16, and 17, which intersect with genes that show a significant excess of DNM (red, turquoise and blue points). The analysis confirms known associations such as *PAFAH1B1* (LIS1), *RAI1,* and *KANSL1* and candidate associations for *MAPK3* and *SIN3A.* Recurrent severe missense of *GABRB3* has been associated with autism and may be relevant to the recurrent 15q11 duplication We note that mutations and deletions of the imprinted genes *SNRPN* (no DNM in our data set) and *UBE3A* (1 LGD and 1 missense DNM in our dataset) are known to cause the core phenotype of Prader-Willi and Angelman syndromes, respectively, but do not reach significance in this analysis.

Many genomic disorders intersect with a single DNM-enriched gene, confirming a known CNV gene association, including *KANSL1* (Koolen-de Vries)^57^, *PAFAH1B1* aka *LIS1* (Miller-Dieker)^58^, *SHANK3* (Phelan-McDermid)^59^, *RAI1* (Smith-Magenis)^60^, *NSD1* (Sotos)^61^, *WHSC1* (Wolf-Hirschhorn)^62^, *BCL11A* (2p15-16.1 microdeletion)^63^, and *CREBBP* (Rubinstein-Taybi)^64^ (Table 3). In addition, this analysis also highlights genes that have been implicated as candidates by case reports, functional studies, or smaller CNVs (Table 3). Among these, we identify an excess of recurrent missense mutation in *MAPK3* mapping to the 16p11.2 microdeletion/microduplication region associated with autism and ID^65^. Recurrent LGD mutations in *PHF21A,* a gene previously implicated by translocations and focal CNVs^66,67^, map to the Potocki-Shaffer deletion region, while an excess of missense mutations in *KIF1A* correspond to the 2q37 deletion syndrome region^68,69^. Recurrent LGD DNMs in *SIN3A,* a REST and MECP2 interactor, map to the 15q24 deletion region^44,70,71^. *CLTC* has been linked to multiple malformations and DD and is located in the 17q23 deletion region^72^. Genes enriched for recurrent missense DNM, *YWHAG* and *GABRB3,* co-localize to the Williams-Beuren and Prader-Willi deletion/duplication regions, respectively^73–75^.

Finally, we also considered as part of this analysis the 14 regions identified as significant for CNV burden^18^ and identified five candidate intersections (Table 3). These include *SATB2* in the 2q33.1 region ^18,76^; *MEF2C,* which demonstrated focal deletions, functions at cortical synapses and has been independently linked to hyperkinesis and epilepsy^77,78^; *CHD4* in the 12p13 duplication region, which has been linked by both a genome-wide association study and DNM to an ID syndrome^79,80^; *WDFY4* in the 10q11.23 duplication region, which appears to be a novel finding at this time; and, finally, *PAX5* in the 9p13 region, which was previously highlighted in O’Roak et al.^81^.

A second category of CNVs are those that intersect more than a single gene (Table 3). The large 9q34 deletion region, for example, contains two genes with recurrent LGD mutations: the expected Kleefstra syndrome gene *EHMT1*^82^ and a relatively unknown gene, *C9orf142,* implicated in DNA repair (see above)^83^. The region also harbors *GRIN1,* where recurrent missense mutations have been identified in DD^84^. The 2q33.1 region contains several potentially high-impact candidates, including *HECW2,* which has been linked to neurodevelopmental delay, ID and epilepsy by missense mutations^26,85^; *SATB2,*which has been independently identified by focal CNVs^18,76^; *ABI2,* which is a candidate for autosomal recessive ID^86^; and *SF3B1,* which interacts directly with the ID gene *PQBP1^87^.* In the Sotos syndrome region we identify not only the critical gene *NSD1^61^* but also significance in *GPRIN1* by LGD enrichment (denovolyzeR model only). Similarly, in the 2p15-p16.1 deletion region we identify both the primary gene *BCL11A^63^* as well as a second candidate gene in the minimal critical region *PAPOLG^88^,* which is enriched for severe missense DNM. In the 17q23.1q23.2 deletion we identify enrichment in *PPM1D,* which has been directly linked to ID^50^, and *MED13,* which is also part of the minimal region but only marginally significant for *de novo* enrichment. Finally, in the 2q 11.2q13 deletion region we identify both *POU3F3,* which has been linked to ID and dysmorphic features by focal deletions^89^, and *RFX8,* which has limited functional information in the literature.

## DISCUSSION

Exome sequencing of parent–child trios is a particularly powerful tool for the identification of genes, which when disrupted lead to pediatric NDD. While CNV maps historically offered sensitivity by detecting submicroscopic structural variants below the level of karyotype and high rates of recurrence based on the genome duplication architecture, exome sequencing added a new level of specificity allowing the identification of single genes by detection of disruptive *de novo* variants in the entire coding sequence of the human genome. Here, we expand and integrate the data by performing a meta-analysis of DNMs in 10,927 children with autism and NDDs with CNV morbidity data. The use of two DNM models (CH model and denovolyzeR) identifies a high-confidence intersection (n = 164 genes) and a comprehensive union (301 genes), which reach significance by one or both models. The combined analysis leverages the unique strengths of each model, such as the coding sequence chimpanzee–human divergence metrics in the CH model, and high-quality triplet context-based mutation rates in denovolyzeR. Examination of this gene set in the context of general population conservation statistics based on the ExAC population (pLI and missense Z scores from 45,376 individuals excluding neuropsychiatric cases, RVIS from the full 60,706 cases) confirms that this set list is enriched for genes that are constrained in the general population (LGD pLI 2.7e-62, Missense Z p=1.3e-51, RVIS p < 1e-7 to 5e-5) (Figure 1).

While intolerance metrics are useful to enrich for pathogenic genes, our analysis suggests caution in strict application of a specific cutoff or even a single metric. Several known pathogenic genes are borderline by only one intolerance score, while several are poorly constrained by both metrics (e.g. *MECP2* and Rett syndrome, RVIS = 32.4, pLI = 0.66). For example, we identify 22 genes that are intolerant to mutation by pLI but not RVIS (RVIS > 20). Some of these are well-established genes (e.g., *KANSL1* and the Koolen-de Vries syndrome)^57^ and the basis for this discrepancy is unknown but may relate to the fact that part of the gene is duplicated complicating genome-wide analyses of intolerance. Among targets poorly constrained by both metrics, the Bohring-Opitz syndrome gene, *ASXL1,* was recently highlighted for the presence of somatic mosaic variants in the ExAC population (from which both the RVIS and pLI scores are derived)^90^. This suggests challenges in the interpretation of disorders that may be driven by both de *novo* and somatic variation in the context of control exomes with almost no phenotypic detail. The presence of several X-chromosome linked genes in this set strongly suggests that gender will also have to be considered in population controls to account for sex-chromosome-specific differences, and potential female protective effects.

Application of subsampling and forward projection estimates indicates under the current DNM enrichment models that gene discovery based on recurrent LGD or severe missense mutations (MIS30) will soon plateau (Figure 4). Simply increasing sample size in this context may be in itself insufficient as the data suggest we are approaching an “n of 1” problem with respect to new gene discovery. Partitioning patients based on additional phenotypic criteria, sub-selecting genes based on functional pathway enrichment^51^, integration of inherited variation^91^, or targeting a small number of genes in much larger cohorts^10,11^ are all strategies for increasing sensitivity for such classes of mutation. For example, analysis of this set of DNMs in the context of MAGI modules further identified key functional categories, including neurotransmitter/synaptic signaling and JNK/MAPK cascades. This analysis identifies 46 genes with DNMs among the four modules that do not yet reach significance but likely represent functionally important targets in future screens.

In contrast to LGD and MIS30 DNM, the number of genes that will be identified by missense DNM generally has not yet begun to approach an asymptote. Samples sizes are just now beginning to reach the level where signatures of recurrence and missense clustering are being detected for a relatively modest number of genes—most of which are only nominally significant^14,15^. It is interesting that many of the nominally significant genes are distinct from those identified by severe recurrent DNM identifying new sets of patients and new candidate genes for clinical investigation and potential therapeutic targets. We propose that this class of mutation (less severe or clustered missense DNMs) represents the most promising reservoir for future gene discovery. As the number of exomes grows for both ASD and DD, the maintenance and curation of de novo databases will be especially important in this regard^19^ as well as the development of more sophisticated missense clustering algorithms^15^.

Previous studies have implicated larger CNVs and increased mutation burden with more severe phenotypic outcomes and here we observe that the majority of DNM in the 301 genes originates from ID/DD cases (>3:1). Importantly, DNM-enriched genes significantly overlap known pathogenic CNV regions (p = 0.0005, LR+ 2.53 [95% CI 1.43 - 1.91]) supporting a common genetic etiology. These specific targets have offered both independent confirmation of existing single-gene associations (e.g., *KANSL1* in the 17q21.31 region), additional support for candidate genes *(MAPK3* in the 16p11.2 region), and further support for potentially oligogenic effects with multiple compelling candidate genes *(HECW2, SATB2, ABI2,* and *SF3B1* in the 2q33.1 deletion region). This is consistent with recent findings suggesting a role for multiple noncoding mutations in putative regulatory sequence in ASD^92,93^, a feature that cannot be assayed by exome datasets. Among the genes with recurrent mutation and CNV intersection, the *MAPK3* is particularly interesting with respect to the chromosome 16p11.2 microduplication. Several functional studies on 16p11.2 deletion and duplication mice as well as *Drosophila* models have suggested that *MAPK3* is a key regulator of the syndrome: being downstream of other ASD target genes; involved in axon targeting and regulation of cortical cytoarchitecture; and being the most topologically important gene in the region by protein-protein interactions^94–96^. Our analysis builds on these studies by providing evidence of recurrent missense mutation enrichment in human NDDs.

Finally, it is interesting that the 301 genes we highlight in this meta-analysis are enriched for expression in the D1+ and D2+ medium spiny neurons of the striatum (Figure 4). Previously, Dougherty and colleagues highlighted this particular brain region based on a survey of genes reported as autism candidate risk genes^56^. Using a CSEA, we now extend this observation to NDD genes enriched for recurrent DNM. Remarkably, a similar enrichment was recently reported in autistic individuals with multiple DNMs in coding and putative noncoding regulatory DNA^93^. We also observe a similar signature for genes where nominal significance has been observed for clustered DNMs. While many of the genes enriched for D1+ and D2+ expression are not exclusive to the striatum and are more broadly expressed (as demonstrated by the enrichment signal at the lowest specificity threshold), the striatum has been implicated in ID and autism pathology by numerous studies^97–106^. The striatum is particularly compelling as it has been linked to repetitive behaviors^97^ core to the autism phenotype and also to genes known to be involved in DD, including *CHD8, SHANK3, FOXP2,* and *KCNA4*^98,99,102,105,106^. While the striatum is most strongly linked to autism core phenotypes, our observation of enrichment in a more general DD cohort suggests that, while the general bias of cortex genes to ID and striatum genes to ASD^101^ still holds, the diverse expression patterns of genes across the brain at complex developmental time points may have substantial functional overlap among subtypes of NDDs that will require deep phenotyping and imaging to tease apart.

In conclusion, the 301 genes that show evidence for recurrent DNM represent a starting point for further functional and phenotypic investigations. These genes demonstrate strong conservation, refine pathogenic CNVs, define distinct functional pathways, and support the role of striatal networks in the pathogenicity of both ASD and ID/DD. In addition, we identify new candidates by clustered missense mutations, many of which are enriched in synaptic function but appear distinct from those genes where recurrent DNM are observed. Strikingly, the majority of the genes identified in this study present with DNMs in both ASD and ID/DD. While we expect a degree of diagnostic overlap^20,21^, our results support a common genetic etiology among broad neurodevelopmental phenotypes. These genes are candidates for a genotype-first paradigm^107^, where downstream follow-up of patients with the same *de novo* disrupted gene is likely to provide additional insight into unique phenotypic features associated with these different genetic subtypes^108^ and additional support for their role in NDD.

## ACKNOWLEDGEMENTS

We wish to thank Tychele Turner Joseph Dougherty, and Tonia Brown for helpful discussion and edits. E.E.E. is an investigator of the Howard Hughes Medical Institute.

## METHODS

### Data set

We selected *de novo* variation from 10,927 cases with neurodevelopmental diagnoses of ASD or ID/DD compiled in the denovo-db v.1.5 release^19^. All variants were annotated to RefSeq transcripts using SNPEFF and collapsing to the most severe variant across isoforms. Variants were further binned into LGD (stop loss/gain, splice, and frameshift), missense, or synonymous categories for analysis.

### Statistical analysis

Enrichment of *de novo* LGD and missense variation per gene was calculated using two statistical models. The CH model as previously described^11^ was run using the default setting and assuming a baseline rate of 1.8 de novo variants per individual. In addition, we ran a recently published modified version^39^ that separately tests for enrichment of variants with CADD v.1.3 scores^38^ of 30 and higher, which are predicted to be the most damaging of missense variation. Similarly denovolyzeR^12^ was run using default settings. Each test (LGD, missense, MIS30) was individually adjusted to a q-value by the Benjamini–Hochberg procedure based on the number of genes in the model (exome wide) and genes with q < 0.1 and a DNM count of 2 or more were considered for the union set. Wherever necessary, gene symbols were adjusted to match those used in the individual models (CH model, denovolyzeR, CSEA, CLUMP). Detection of excess missense clustering in genes was performed using the CLUMP tool as previously described^1314^. Briefly, we implemented the permutation (-z 1000) and minimum mutation options (-m 2) and calculated a p-value based on the null distribution of case and control CLUMP score differences. To enable exact transcript comparisons between cases and controls, all variants were re-annotated with CRAVAT^109^ resulting in annotation of 9,772 de novo missense variants in cases. The control set included private missense mutations in individuals from ExAC without neuropsychiatric disorders (n = 45,376; 1,466,439 mutations). All genes with at least two mutations in cases and controls were tested (n = 1930). For each gene we initially performed 1,000 simulations and in cases where nominal significance was reached we performed 1,000,000 simulations and calculated an empirical p-value^110^. Functional enrichment was examined using the CSEA and TSEA tools^56^. Identification of clustered gene modules was performed using the MAGI (merging affected genes into integrated networks)^51^ enrichment tool with default settings and incorporating co-expression and physical interaction data from geneMANIA^111^. Prediction of future LGD and missense variation discovery rates was determined by sampling (with replacement) populations of 100 to 10,900 cases 10,000 times each and calculating DNM statistics using the CH model. The number of genes with two or more mutations and a q-value ≤ 0.1 were then enumerated for each simulation and linear as well as logistic growth models were fit to each curve with the best model being chosen by BIC. Model fits and confidence bounds were performed using the base stats and propagate (https://CRAN.R-project.org/package=propagate) packages in the R statistical language.

---

**Table S1:**
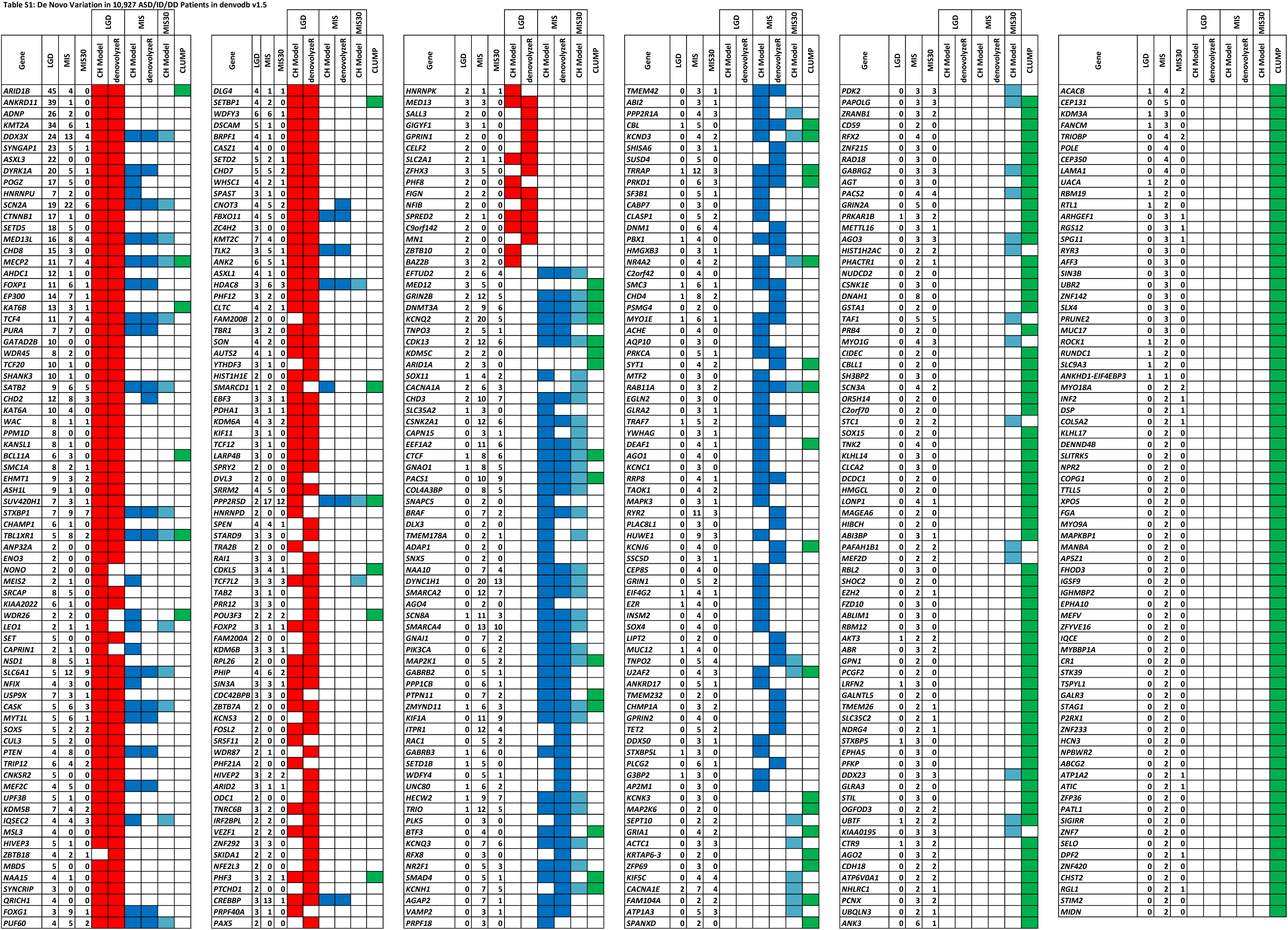
De Novo Variation in 10,927 ASD/ID/DD Patients in denvodb v1.5

